# Revising mtDNA haplotypes of the ancient Hungarian conquerors with next generation sequencing

**DOI:** 10.1101/092239

**Authors:** Endre Neparáczki, Klaudia Kocsy, Gábor Endre Tóth, Zoltán Maróti, Tibor Kalmár, Péter Bihari, István Nagy, György Pálfi, Erika Molnár, István Raskó, Tibor Török

## Abstract

As part of the effort to create a high resolution representative sequence database of the medieval Hungarian conquerors we have resequenced the entire mtDNA genome of 24 published ancient samples with Next Generation Sequencing, whose haplotypes had been previously determined with traditional PCR based methods. We show that PCR based methods are prone to erroneous haplotype or haplogroup determination due to ambiguous sequence reads, and many of the resequenced samples had been classified inaccurately. The SNaPshot method applied with published ancient DNA authenticity criteria is the most straightforward and cheapest PCR based approach for testing a large number of coding region SNP-s, which greatly facilitates correct haplogroup determination.

## Introduction

Comparing ancient DNA (aDNA) sequences extracted from well dated archaeological remains from different periods and locations provide crucial information about past human population history (reviewed in (Pickrell and Reich, 2014). Phylogeographic inferences are drawn from phylogenetic and population genetic analyses of sequence variations, the quality of which can be biased by data quantity and quality. Nowadays Next Generation Sequencing technology (NGS) provides a growing number of high quality aDNA sequence data, but until recently the majority of aDNA studies have been restricted to short fragments from the hypervariable region-1 (HVR-I) of the mitochondrial DNA (mtDNA) genome, using PCR based methods. PCR based methods are very sensitive for contamination, as low amounts of exogenous DNA can easily dominate PCR products resulting in the recovery of irrelevant sequences (Richards et al., 1995) (Malmström et al., 2005) (Pilli et al., 2013) (Heupink et al., 2016). As a result, in spite of the applied authenticity criteria (Knapp et al., 2012), many of the published databases may contain unreliable sequences, which distort statistical analyses. This problem is especially relevant for many of the ancient populations, from which only PCR based HVR data are available.

Recently several aDNA studies were published aiming to shed light on the origin of ancient Hungarians, two of these (Tömöry et al., 2007) (Csősz et al., 2016) applied restriction fragment length polymorphism (RFLP) to identify 11 and 14 haplogroup (Hg) specific coding region SNP-s in addition to HVR sequencing, while another study (Neparáczki et al., 2016) tested 22 coding region SNP-s with multiplex PCR and GenoCoRe22 assay described in (Haak et al., 2010).

Using the NGS method combined with hybridization enrichment, we have sequenced the entire mtDNA genome of 9 samples from the (Tömöry et al., 2007) study, and 15 samples from the (Neparáczki et al., 2016) study, so we could compare the reliability of two different traditional approaches.

## Materials and Methods

### Archaeological samples

Bone samples from the Hungarian conquest period used in the study of (Tömöry et al., 2007) are carefully maintained in the anthropological collection at the Department of Biological Anthropology, University of Szeged, Hungary, so we could unambiguously identify and resample these remains. Bone powder remains of samples from the study of (Neparáczki et al., 2016), were saved in the Department of Genetics, University of Szeged, and were reused to build NGS sequencing libraries.

### DNA extraction

Ancient DNA work was performed in the specialized ancient DNA (aDNA) facilities of the Department of Genetics, University of Szeged, Hungary with strict clean-room conditions. 100 mg bone powder from tooth roots or petrous bones was predigested in 1 ml 0,5 M EDTA 100 μg/ml Proteinase K for 30 minutes at 48°C, to increase the proportion of endogenous DNA (Damgaard et al., 2015), then DNA solubilisation was done overnight, in 1 ml extraction buffer containing 0.45 M EDTA, 250 μg/ml Proteinase K, 1% Triton X-100, and 50 mM DTT. DNA was bound to silica (Rohland and Hofreiter, 2007) adding 6 ml binding buffer (5,83 M GuHCl, 105 mM NaOAc, 46,8% isopropanol, 0,06% Tween-20 and 150 μl silica suspension to the 1 ml extract, and the pH was adjusted between 4–6 with HCl. After 3 hours binding at room temperature silica was pelleted, and washed twice with 80% ethanol, then DNA was eluted in 100 μl TE buffer.

### NGS library construction

First 50 μl DNA extract was subjected to partial uracil-DNA-glycosylase (UDG) treatment followed by blunt end repair, as described in (Rohland et al., 2015a). DNA was then purified on MinElute column (Qiagen), and double stranded library was made as described in (Meyer and Kircher, 2010), except that all purifications were done with MinElute columns, and after adapter fill-in libraries were preamplified in 2 x 50 μl reactions containing 800 nM each of IS7 and IS8 primers, 200 μM dNTP mix, 2 mM MgCl_2_, 0,02 U/μl GoTaq G2 Hot Start Polymerase (Promega) and 1X GoTaq buffer, followed by MinElute purification. PCR conditions were 96°C 6 min, 16 cycles of 94 °C 30 sec, 58 °C 30 sec, 72 °C 30 sec, followed by a final extension of 64 °C 10 min. Libraries were eluted from the column in 50 μl 55 °C EB buffer (Qiagen), and concentration was measured with Qubit (Termo Fisher Scientific). Libraries below 5 ng/μl concentration were reamplified in the same reaction for additional 5–12 cycles, depending on concentration, in order to obtain 50 μl preamplified library with a concentration between 10–50 ng/μl.

50 ng preamplified libraries were double indexed according to (Kircher et al., 2012) in a 50 μl PCR reaction containing 1 x KAPA HiFi HotStart ReadyMix (Kapa Biosystems) and 1000 nM each of P5 and P7 indexing primers. PCR conditions were 98 °C 3 min, 6 cycles of 98 °C 20 sec, 66 °C 10 sec, 72 °C 15 sec followed by a final extension of 72 °C 30sec. Indexed libraries were MinElute purified and their concentration was measured with Qubit, and size distribution was checked on Agilent 2200 TapeStation Genomic DNA ScreenTape.

Control libraries without UDG treatment were also made for assessing the presence of aDNA specific damages in the extract, as well as DNA free negative control libraries, to detect possible contamination during handling or present in materials.

### Mitochondrial DNA capture and sequencing

Biotinilated mtDNA baits were prepared from three overlapping long-range PCR products as described in (Maricic et al., 2010), but using the following primer pairs, L14759-H06378, L10870-H14799, L06363-H10888, described in (Haak et al., 2010).

Capture was done according to (Maricic et al., 2010) with the following modifications: Just four blocking oligos, given below were used in 3 μM (each) final concentration: BO1 P5.part1F: AATGATACGGCGACCACCGAGATCTACAC-Phosphate, BO2.P5.part2F ACACTCTTTCCCTACACGACGCTCTTCCGATCT-Phosphate, BO4.P7.part1 R GTGACTGGAGTTCAGACGTGTGCTCTTCCGATCT-Phosphate, BO6.P7.part2 R CAAGCAGAAGACGGCATACGAGAT-Phosphate.

For one capture 300 ng biotinilated bait was used with 30 jal Dynabeads MyOne Streptavidin C1 magnetic beads (Thermo Fisher Scientific). Double indexed libraries of 20 samples (300 ng each) were mixed and concentrated on MinElute columns, then captured together in a 64 μl hybridization reaction. When fewer samples were enriched, we used proportionally smaller amounts of bates. After washing, bead-bound enriched libraries were resuspended in 20 μl water and released from the beads in a 60 μl PCR reaction containing 1 X KAPA HiFi HotStart ReadyMix and 2000 nM each of IS5- IS6 library primers. PCR conditions were: 98 °C 1 min, 10 cycles of 98 °C 20 sec, 60 °C 30 sec, 72 °C 30 sec, followed by a final extension of 72 °C 30 sec. The captured and amplified library mix was purified on MinElute column and eluted in 15 μl EB.

Before sequencing, libraries were quantified with Qubit, and quality checked and Agilent 2200 TapeStation Genomic DNA ScreenTape. Sequencing was done at the SeqOmics Biotechnology Ltd., using MiSeq sequencer with MiSeq Reagent Kit v3 (Illumina, MS-102–3003) generating 2x150bp paired-end sequences.

### Data analysis

The adapters of paired-end reads were trimmed with the cutadapt software (http://dx.doi.org/10.14806/ej.17.1.200) in paired end mode. Read quality was assessed with FastQC (http://www.bioinformatics.babraham.ac.uk/projects/fastqc/). Sequences shorter than 25 nucleotide were removed from this dataset.The resulting analysis-ready reads were mapped to the GRCh37.75 human genome reference sequence using the Burrows Wheeler Aligner (BWA) v0.7.9 software (Li and Durbin, 2009) with the BWA mem algorithm in paired mode and default parameters. Aligning to the GRCh37.75 human reference genome that also contains the mtDNA revised Cambridge Reference Sequence (rCRS, NC_012920.1) (Andrews et al., 1999) helped to avoid the forced false alignment of homologous nuclear mitochondrial sequences (NumtS) to rCRS, though the proportion of NumtS, derived from low copy nuclear genome, is expexted to be orders of magnitudes lower than mtDNA in aDNA libraries. Samtools v1.1 (Li et al., 2009) was used for sorting and indexing BAM files. PCR duplicates were removed with Picard Tools v 1.113 (http://picard.sourceforge.net). Ancient DNA damage patterns were assessed using MapDamage 2.0 (Jónsson et al., 2013), and read quality scores were modified with the rescale option to account for post-mortem damage. Freebayes v1.02 (arXiv: 1207.3907 [q-bio.GN]) was used to identify variants and generate variant call format (VCF) files with the parameters -q 10 (exclude nucleotids with <10 phred quality) and -P 0.5 (exclude very low probability variants). Each variant call was also inspected manually. From VCF files FASTA format was generated with the Genom Analysis Tool Kit (GATK v3.5) FastaAlternateReferenceMaker walker.

## Results and Discussion

### NGS sequencing

We have sequenced 24 complete mtDNA genomes of the ancient Hungarians with multiple coverage (Table 2) without gaps and determined the haplotypes of the individuals (Table 1 and Supplementary Table1). For two samples we have replicated the experiments from two independent extracts, one from bone another from tooth derived from the same individual, and in each case received identical sequence reads. UDG treated and non UDG treated libraries derived from the same extract also gave the same sequence reads. MapDamage profile of our partial UDG treated and control non treated library molecules displayed typical aDNA damage distribution (Supplementary Figure 1), as described in (Rohland et al., 2015b). MapDamage computed proportions of sequence reads with aDNA specific C-to-T and G-to-A transitions at the ends of molecules which remained after partial UDG treatment are shown in Table 2. The average length of the obtained mtDNA fragments ranged from 56 to 85 bp (Table 2), an expected size range for aDNA (Sawyer et al., 2012). These data indicated that the majority of sequences were derived from endogenous DNA molecules. Then we have estimated the percentage of possible contaminating molecules (Table 2) with a similar logic as in (Fu et al., 2013), by calculating the proportion of reads which did not correspond with the diagnostic positions of the consensus sequence given in Supplemetary Table 1, which revealed very low contamination levels. Phylogenetic analyses (HaploGrep 2, (Weissensteiner et al., 2016) of all consensus sequences resulted unambigous classifications without contradictory positions. Consensus sequences were submitted to NCBI GenBank under Accession No: KY083702-KY083725.

**Table 1.**
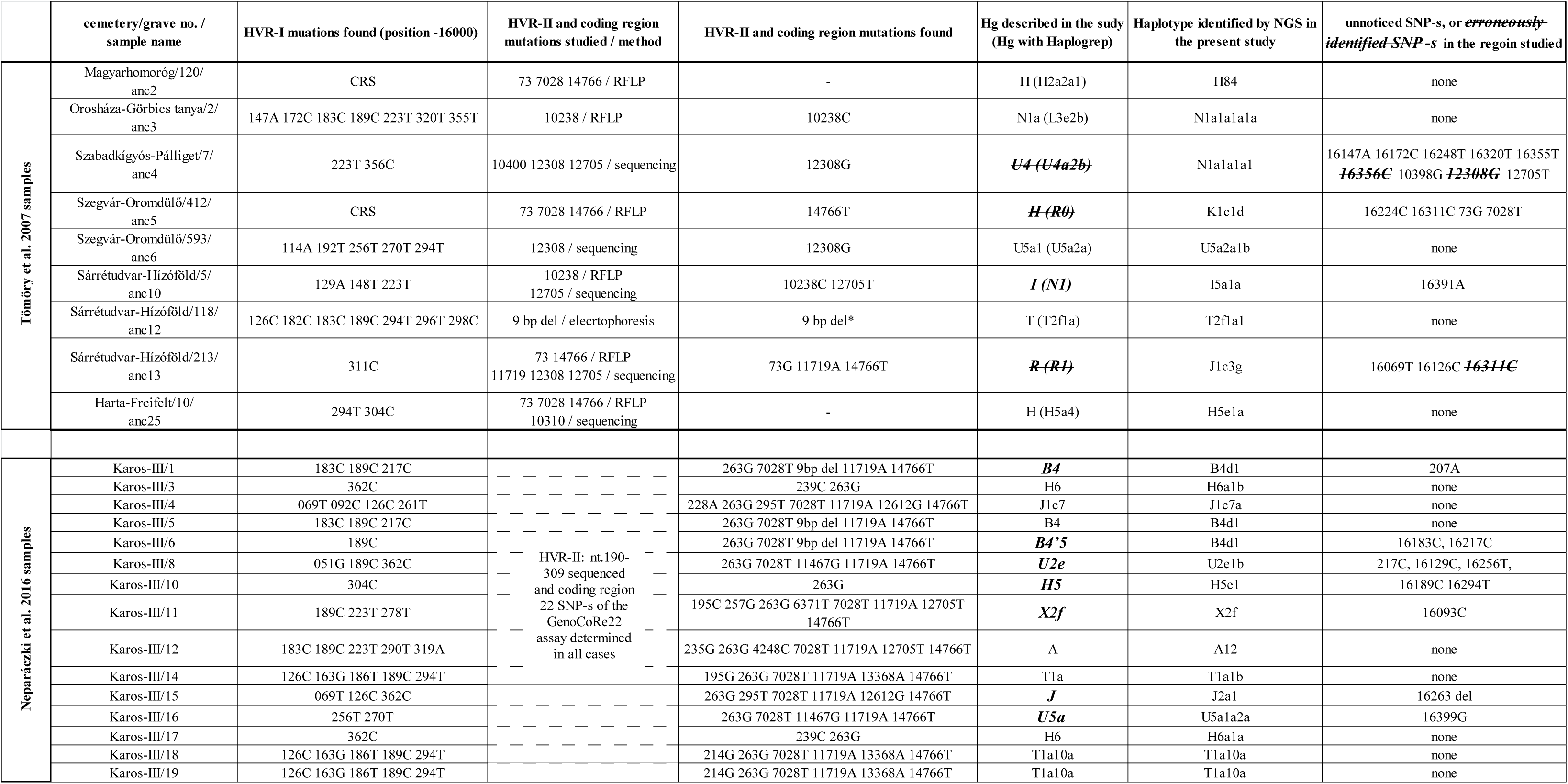
Comparison of Haplogroups identified with different PCR based methods and NGS. Hg-s determined incorrectly with PCR methods are labelled with enlarged bold italic and lined through, while correct Hg-s with incorrect haplotypes are labelled with enlarged bold italic. Haplogroups and haplotypes were determined with the HaploGrep 2 version 2.1.0 (Weissensteiner et al., 2016) based on Phylotree 17 (van Oven, 2015) from the available SNP-s. For the (Tömöry et al., 2007) samples HaploGrep assignment, based on their identified SNP positions is given in parenthesis. *data from Ph.D thesis of Tömöry 2008.

**Table 2.** Details of NGS data for each samples. Data shown are all from UDG treated libraries.

| cemetery/grave no.<br>/sample name | sample<br>source | total no.<br>of reads | no. of<br>reads<br>mapped on<br>rCRS | no. of<br>unique<br>mapped<br>reads | average<br>fragment<br>length | (%) of<br>nucleotides<br>above 10x<br>coverage | estimated<br>contaminat<br>ion (%) | (%) G to A<br>misincorp. at<br>3' end<br>(MapDamage) | (%) C to T<br>misincorp. at<br>5' end<br>(MapDamage) |
| --- | --- | --- | --- | --- | --- | --- | --- | --- | --- |
| Magyarhomoróg/120<br>/anc2 | tooth | 26152 | 12519 | 1656 | 55,97 | 89,62 | 0,00 | 6,11 | 7,81 |
| Orosháza-Görbics<br>tanya/2/ anc3 | femur | 85178 | 50555 | 3444 | 79,37 | 99,89 | 0,43 | 8,13 | 8,89 |
| Szabadkigyós-Pálliget/7/anc4 | tooth | 53176 | 27002 | 2854 | 63,85 | 97,46 | 0,45 | 8,19 | 9,22 |
| Szegvár-Oromdűlő/412/anc5 | tooth | 47760 | 23426 | 6150 | 60,07 | 99,95 | 0,75 | 5,49 | 6,40 |
| Szegvár-Oromdűlő/593/anc6 | tooth | 66798 | 35260 | 3841 | 58,09 | 98,64 | 2,08 | 6,47 | 8,70 |
| Sárrétudvari-Hízóföld/5/anc10 | tooth | 17632 | 6664 | 2205 | 69,94 | 95,11 | 0,00 | 7,40 | 10,29 |
| Sárrétudvari-Hízóföld/118/anc12 | tooth | 36214 | 14708 | 3394 | 63,96 | 98,81 | 0,36 | 8,75 | 11,53 |
| Sárrétudvari-Hízóföld/213/anc13 | tooth | 42326 | 20383 | 4997 | 62,44 | 95,01 | 0,83 | 9,01 | 10,22 |
| Harta-Freifelt/10/anc25 | tooth | 135472 | 66169 | 4803 | 80,75 | 99,93 | 1,75 | 6,00 | 7,10 |
| Karos-III/1 | femur | 30830 | 5968 | 2414 | 81,44 | 98,07 | 0,00 | 13,09 | 11,48 |
| Karos-III/3 | femur | 58982 | 12858 | 3839 | 76,84 | 98,91 | 1,68 | 12,51 | 10,66 |
| Karos-III/4 | femur | 82194 | 22930 | 4862 | 68,71 | 99,98 | 0,33 | 12,93 | 11,98 |
| Karos-III/5 | metatarsus | 394292 | 23740 | 10299 | 85,35 | 99,96 | 3,44 | 8,45 | 6,84 |
| Karos-III/6 | femur | 41054 | 3863 | 955 | 70,68 | 87,24 | 0,00 | 6,71 | 6,75 |
| Karos-III/8 | femur | 75724 | 26747 | 5938 | 64,26 | 99,67 | 1,79 | 11,37 | 11,08 |
| Karos-III/10 | femur | 67416 | 6371 | 1768 | 68,22 | 89,61 | 0,00 | 9,89 | 9,04 |
| Karos-III/11 | femur | 346124 | 4496 | 1569 | 79,62 | 93,91 | 1,32 | 12,78 | 11,42 |
| Karos-III/12 | femur | 52738 | 5843 | 2406 | 70,76 | 95,80 | 3,96 | 12,40 | 11,19 |
| Karos-III/14 | femur | 61346 | 16134 | 4532 | 69,29 | 99,69 | 1,82 | 14,53 | 14,13 |
| Karos-III/15 | femur | 422796 | 80431 | 13248 | 81,03 | 99,98 | 0,29 | 11,02 | 9,95 |
| Karos-III/16 | femur | 90950 | 23233 | 2851 | 75,65 | 99,14 | 1,34 | 11,34 | 11,12 |
| Karos-III/17 | femur | 43330 | 2626 | 1191 | 79,64 | 87,55 | 0,00 | 10,31 | 9,91 |
| Karos-III/18 | femur | 9184 | 3208 | 1596 | 68,48 | 90,49 | 0,00 | 15,42 | 12,11 |
| Karos-III/19 | tooth | 59102 | 30135 | 3244 | 69,07 | 98,78 | 0,00 | 6,39 | 6,55 |

In NGS sequence reads typical aDNA sequence alterations, present in individual molecules, are disclosed and excluded by averaging multiple reads. Moreover aDNA specific sequence alterations, primarily C-T and G-A transitions accumulating at the end of molecules, serve as markers to distinguish ancient molecules from contaminating modern DNA. Therefore NGS eliminates most sequencing uncertainties inherent in PCR based methods (reviewed in (Rizzi et al., 2012), resulting in very reliable sequence reads. So we could use our NGS data to reevaluate and compare previous haplotyping strategies used in (Tömöry et al., 2007), (Csősz et al., 2016) and (Neparáczki et al., 2016). For this end, from our NGS data, we collected all SNP-s within the HVR stretches and coding region positions, which had been examined in (Tömöry et al., 2007) and (Neparáczki et al., 2016), then contrasted these with the original dataset (Table 1).

### Contrasting NGS and PCR based sequence data

We found that in (Tömöry et al., 2007) haplotypes of 5 out of 9 samples were determined correctly, while in one sample haplogroup was correct with inaccurate haplotype, and in 3 samples NGS detected entirely different haplogroups. In the 15 samples of (Neparáczki et al., 2016) the same haplogroups were assigned from NGS data in all cases, however only 8 haplotypes proved to be correct. In both studies the majority of deviations originated from undetected SNP-s in sequencing reactions of PCR fragments, but (Tömöry et al., 2007) also identified 3 SNP-s erroneously (lined through nucleotide positions in Table 1). These results indicate that haplotypes from both studies were rather unreliable, but haplogroup classification with the approach of (Neparáczki et al., 2016) is more trustworthy than with approach used in (Tömöry et al., 2007).

As multicopy mtDNA is much better best preserved in archaeological remains than low copy nuclear DNA, most ancient sequences are derived from mitochondria (Hofreiter et al., 2001). Within mtDNA, the most polymorphic HVR control region contains outstanding phylogenetic information, therefore HVR sequencing has been the primary method of choice for mtDNA hapolotyping. However HVR polymorphisms have a limited reliability for haplogroup determination, therefore in addition several informative coding region SNP-s (CR-SNP) were selected to unambiguously define haplogroups (Behar et al., 2007). At the beginning individual CR-SNP-s were determined with RFLP (Brown, 1980) or direct sequencing of PCR clones, but soon multiplex PCR combined with the SNaPshot technique (Salas et al., 2005) offered a more straightforward solution for identifying multiple SNP-s simultaneously. Latter method was soon adapted in the ancient DNA field (Bouakaze et al., 2007), (Haak et al., 2010).

Determining individual CR-SNP-s separately is very time consuming and expensive, so it is tempting to test just those CR-SNP-s which are in line with HVR-I data. This is exactly what we read in (Tömöry et al., 2007): *“In cases when haplogroup categorization was not possible on the basis of HVSI motifs alone, analysis of the diagnostic polymorphic sites in the HVSII region and mtDNA coding region was also performed.”* A major problem with this approach is the ambiguity of sequence reads derived from aDNA PCR clones, as amplification typically starts from a mixture of endogenous and contaminating human DNA molecules (Malmström et al., 2005). Erroneous HVR reading will lead to inappropriate CR-SNP selection, and in case of dubious CR-SNP results, false Hg classification. This is the most probable explanation of the 3 incorrectly defined haplogroups in (Tömöry et al., 2007), (Table 1). A major advantage of the GonoCore22 SNaPshot assay is that all Hg specific CR-SNP-s are examined irrespectively of HVR reads. The 22 CR-SNP alleles independently define a certain Hg, which must correspond with that based on HVR sequence. As both HVR and CR-SNP reads may give ambiguous results, this approach provides a double control for correct Hg designation, but is not immune against incorrect HVR haplotype reads. This is the explanation of correct Hg-s and erroneous haplotypes in (Neparáczki et al., 2016), (Table 1).

The problem of ambiguous aDNA sequence reads is demonstrated on Fig. 1. In (Neparáczki et al., 2016) consequently the higher peaks were taken into account, which also matched with the GenoCoRe22 data. However in position 16399 the correct nucleotide is defined by the neglected lower peak (G instead of A, see Table 1.), which resulted in incorrect haplotyping. In contrast in the neighboring double peak (16403 in Fig. 1), the correct nucleotide is defined by the selected higher peak.

**Figure 1.**
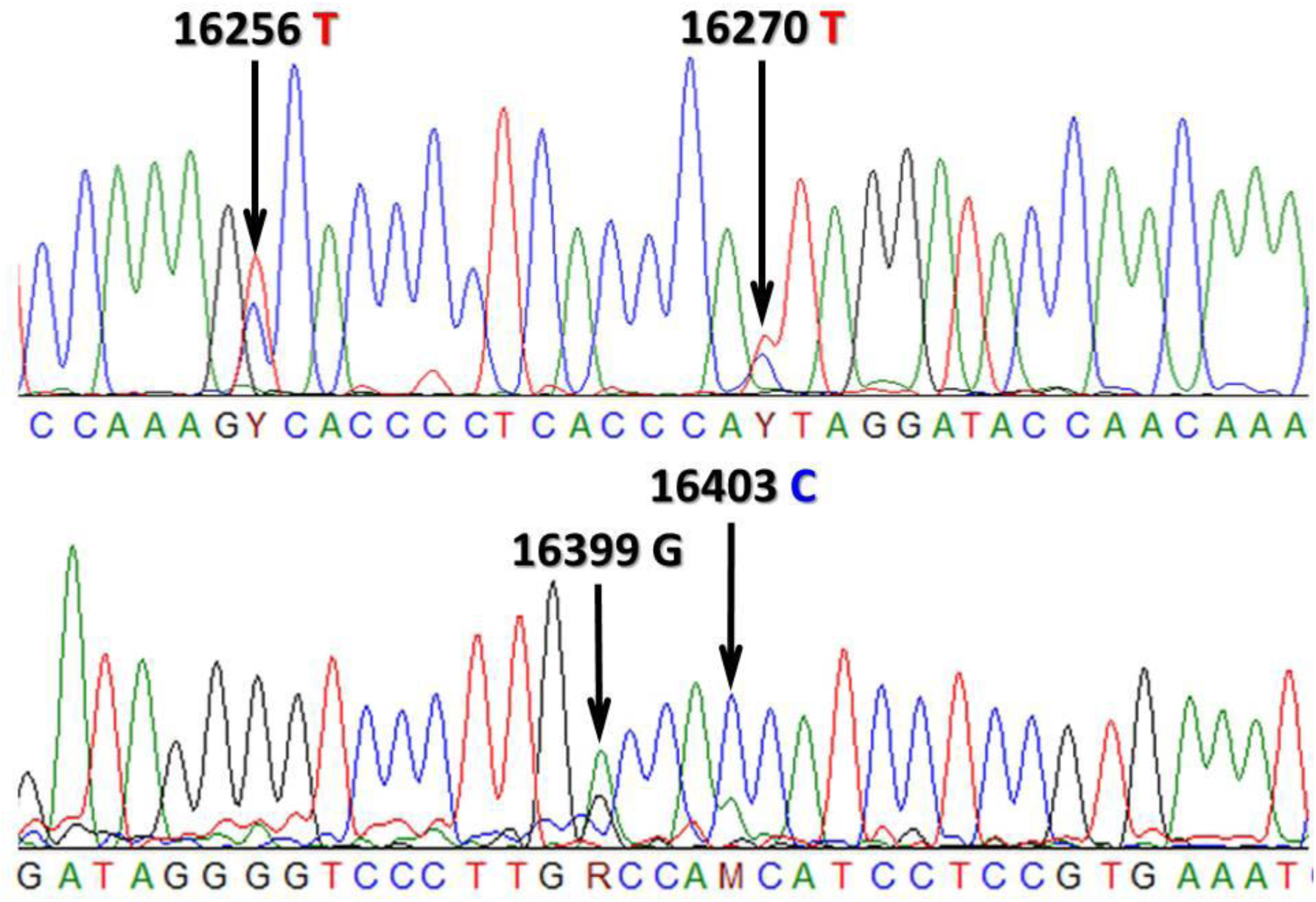
Chromatogram of two HVR-I sequence fragments of the Karos-III/16 sample from (Neparáczki et al., 2016). Arrows label double peaks, correct reads according to NGS data are listed above the arrows.

Coding region SNP testing with either RFLP, sequencing or SNaPshot method also suffers from the same problem as demonstrated on Fig. 2. After multiplex PCR amplification of 22 mtDNA fragments two separate Single Base Extension (SBE) reactions are performed, and each reveals 11 Hg defining alleles. Both independent SBE reactions shown in Fig. 2 contain several double peaks, and one of each must have derived from contamination. Some of these can be excluded from repeated SNaPshot reactions, for example the lower electropherogram excludes the ancestral *preHV* allele, since it has a single peak (T) in this position. If such exclusion is not possible, the higher peaks are preferably chosen, as the blue peak (G) for Hg *B* and the green (A) for Hg *N* on Fig. 2. These decisions however must be handled with caution, therefore the presence of the *B* Hg defining 9 bp deletion also had been confirmed in (Neparáczki et al., 2016), with singleplex PCR and agarose gelelectrophoresis. In other cases phylogenetic relations are taken into account (Cooper and Poinar, 2000), for example if the *preHV* allele is derived the *HV* allele must also be derived, this is why we have considered the lower peak (A) for *HV* in Fig. 2 (Neparáczki et al., 2016). The summary of repeated SNaPshot reactions considered together with multiple HVR sequence reads warrants trustable Hg classification.

**Figure 2.**
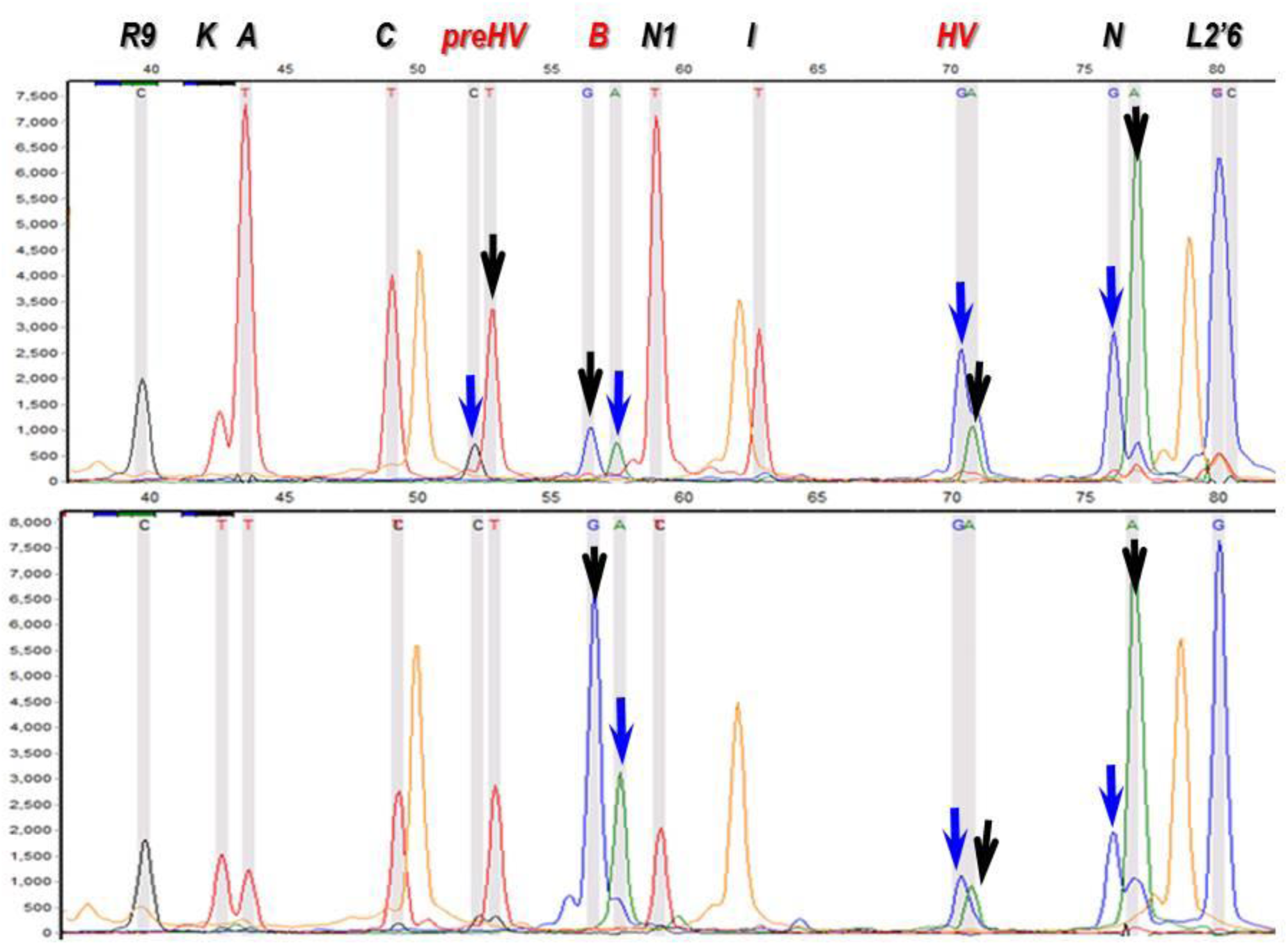
Electropherograms of two SNaPshot SBE-II reactions from two extracts of the same Karos-III/6 sample (Neparáczki et al., 2016). Characters at the top indicate Hg-s defined by the corresponding peaks. Black characters indicate peaks defining the ancestral allele, read characters indicate peaks defining the derived allele. Arrows point at double peaks. As each dye has a different influence on DNA mobility, positions of identical fragments with different dyes are not the same. Black arrows point at peaks taken into account, while blue arrows indicate neglected peaks, considered to have been derived from contamination. Orange peaks are size standards (GeneScan-120 LIZ, Applied Biosystems).

The studied conqueror samples were excavated between the 1930-90s, and had been handled by a large number of researchers, many with untraceable identity. It follows that these samples were inevitably contaminated during sample collection and storage. (Tömöry et al., 2007) collected samples from a large number of cemeteries, and published the ones with best DNA preservation. In spite of careful sampling their available method was error prone. (Neparáczki et al., 2016) aimed at characterizing an entire cemetery which limited the ability of sample selection, so in spite of the more reliable method their haplotype determination proved error prone. The lesson from this study is that PCR based haplotypes need to be handled cautiously, which has been well known in the aDNA field (Handt et al., 1994), (Richards et al., 1995), (Gilbert et al., 2005), (Sampietro et al., 2006). It also follows that incorrect haplotypes particularly distort sequence based statistical analysis, like Fst statistics or shared haplotype analysis applied in (Tömöry et al., 2007) and (Csősz et al., 2016). The accumulation of authentical NGS ancient DNA sequence data in databases will greatly facilitate reliable population genetic studies.

## Acknowledgement

The generous support of Avicenna Foundation grant no. GF/JSZF/814/9/2015 to I.R. and encouragement of professor Miklos Maroth is highly appreciated. This research was also supported in part by OTKA NN 78696.

## Supplementary Figures

**Supplemetary Figure 1.**
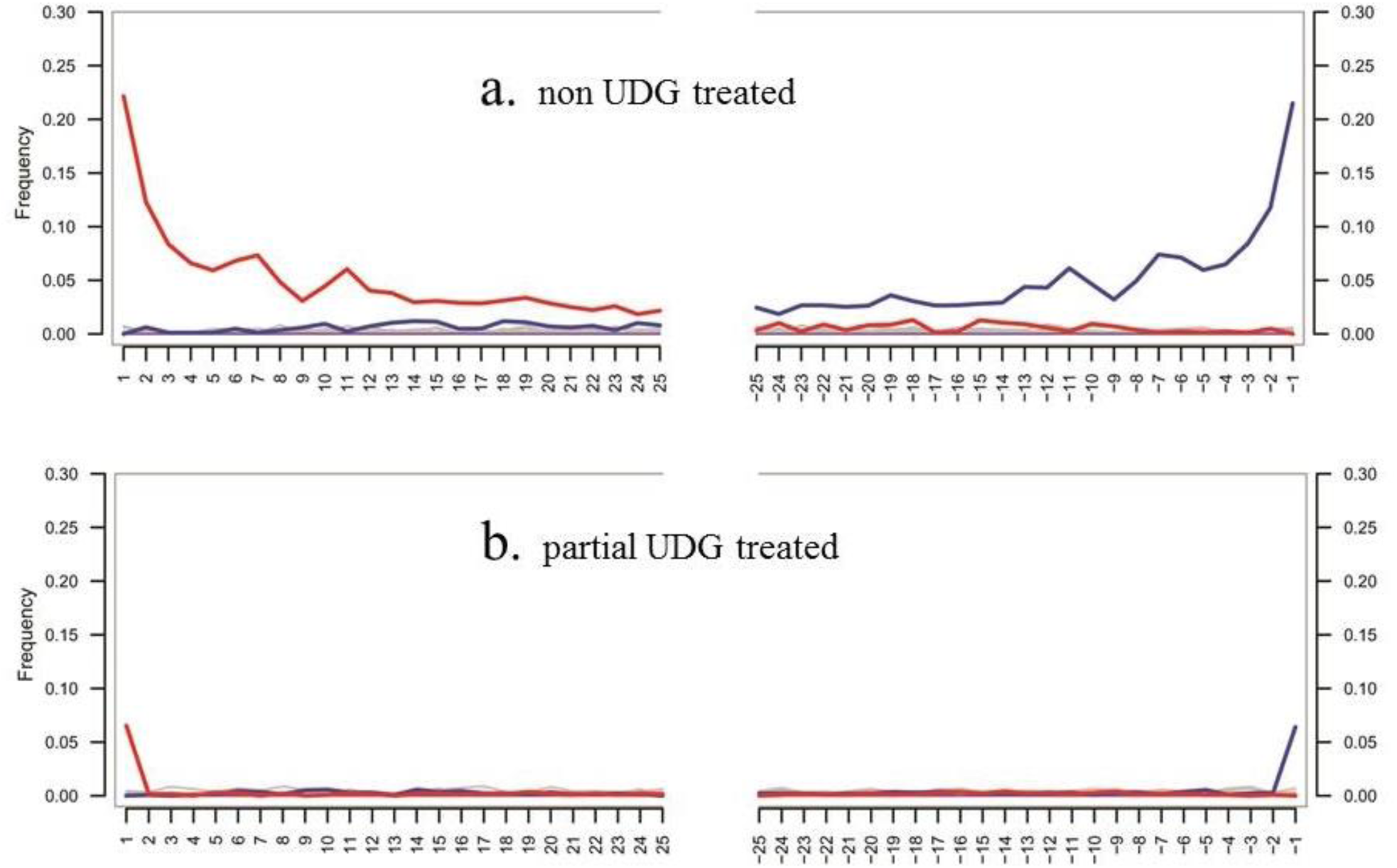
Damage patterns of libraries generated by MapDamage 2.0 20. a. non UDG treated library shownig C to T (and complementary G to A) misincorporations at the 5’ and 3’ termini of the last 25 nucleotides. b. Damage pattern of partial UDG treated library derived from the same extract. As expected the nontreated library contains much higher rate of transitions, most of which was removed by partial UDG treatment. Only data from one extract are shown, as all libraries displayed similar pattern.

## Supplementary Table 1

Mitochondrial sequence haplotypes of the 24 ancient samples. SNPs are provided against rCRS. Following the recommendations in ^36^, we excluded common indels (hotspots) at nucleotide positions: 309.1C(C), 315.1C, 523-524del (or 522-523del), 3106del, 16182C, 16183C, 16193.1C(C), 16519C.

